# Gene flow erodes spatial genomic clines while selection preserves adaptive loci over hundreds of generations

**DOI:** 10.1101/2025.11.28.691231

**Authors:** Vitória Horvath Miranda, Tiago da Silva Ribeiro, Joel M. Alves, Walter Eanes, Evgeny Brud, Krishna Veeramah, John E. Pool, Murillo F. Rodrigues, Rodrigo Cogni

**Affiliations:** Departamento de Ecologia, Instituto de Biociências, Universidade de São Paulo; The Palaeogenomics & Bio-Archaeology Research Network, Research Laboratory for Archaeology and History of Art, University of Oxford; CIBIO, Centro de Investigação em Biodiversidade e Recursos Genéticos, InBIO Laboratório Associado, Universidade do Porto; BIOPOLIS Program in Genomics, Biodiversity and Land Planning, CIBIO; Department of Ecology & Evolution, Stony Brook University; Laboratory of Genetics, University of Wisconsin - Madison; Oregon National Primate Research Center, Oregon Health & Science University

## Abstract

Spatial genetic clines suggest local adaptation, but whether they persist or erode over evolutionary time remains largely unknown. Here, we use a unique genomic time series collected over several decades to quantify how gene flow and selection jointly shape spatial genomic structure. We analyzed pooled whole-genome sequencing from 16 natural populations of *Drosophila melanogaster* sampled along the North American east coast between 1997 and 2023, spanning hundreds of generations. We find pronounced homogenization of populations over time, accompanied by a genome-wide loss of clinal variants. Polymorphisms that remained clinal were less steep on chromosome 3R, reflecting declines in the frequency and clinality of major chromosomal inversions. Genomic context modulates this process: clinal SNPs are preferentially lost in regions of high recombination, whereas remaining clines are increasingly concentrated in coding and splice-associated sites. Despite widespread erosion of spatial structure, we detect signatures of selection shared across space at detoxification loci, including cytochrome P450 genes linked to insecticide resistance. Together, our results show widespread homogenization of neutral spatial variation consistent with gene flow, while spatially varying selection has preserved a subset of functionally important loci.

## Introduction

Comparing historical and contemporary natural populations may illuminate evolutionary changes driven by environmental shifts over time, including those caused by climate change. It also helps us investigate changes caused by demographic processes, such as gene flow and admixture (Clark et al. 2023). Sampling across an environmental gradient adds even more information, and it enables the study of the processes (e.g., selection, isolation-by-distance, admixture) driving genetic differentiation across that gradient. Combining both temporal and spatial dimensions is thus especially powerful for disentangling evolutionary processes (Merilä and Hendry 2014), but most evolutionary studies rely on data from a single time point or location.

Continuous changes in quantitative phenotypes or in allele frequencies along a species range, also called clines, are often regarded as results of local adaptation. Clines have been used to detect selection in many species, such as the malaria vector (*Anopheles gambiae*) (Cheng et al. 2012), the Atlantic salmon (*Salmo salar*) (Vincent et al. 2013), and in barrelclover (*Medicago truncatula* - Fabaceae) (Yoder et al. 2014). However, in many cases, clines are not the product of adaptation to spatially varying environments, but rather the result of demographic processes, such as range expansion (Schäfer et al. 2018), secondary contact between separate populations (Chávez-Galarza et al. 2015), or limited dispersal (Sotka et al. 2004). The investigation of whether clinal patterns reflect local adaptation can greatly benefit from temporal sampling, which requires either ancient DNA or collections made decades apart on organisms with short generation times. Luckily, as with most other patterns, clines have been best studied in model species such as *Arabidopsis* (Lasky et al. 2012; Debieu et al. 2013; Gamba et al. 2024) and *Drosophila* (Klepsatel et al. 2014; Machado et al. 2016; Rajpurohit et al. 2017), which have short generation times that facilitate temporal sampling, as multiple generations occur within each year.

*Drosophila melanogaster* is a particularly good model to investigate evolutionary origins and maintenance of clines. In addition to its status as a model organism with a small, well-annotated genome, phenotypic clines in *D. melanogaster* have been studied since at least 1975 (Adrion et al. 2015). Phenotypic clines in *D. melanogaster* encompass many traits such as body size (James et al. 1995; Zwaan et al. 2000), diapause incidence (Schmidt et al. 2005), and heat and desiccation resistance (Schmidt and Paaby 2008; Rajpurohit et al. 2018). Clines in *D. melanogaster* occur on different continents (Adrion et al. 2015) and are especially well-described along the latitudinal gradient of North America and Australia (Hoffmann and Weeks 2007; Kolaczkowski et al. 2011; Fabian et al. 2012; Adrion et al. 2019).

*D. melanogaster* is native to the African continent (Mansourian et al. 2018) and only recently, about 160 years ago, colonized North America and Australia (Keller 2007). It is possible that at least two distinct *D. melanogaster* populations, each with different genetic backgrounds, colonized North America. One source may have been from Europe, arriving in New York around 1875, and spreading rapidly across the East Coast (Keller 2007). The other source may have reached the Caribbean islands and expanded northward, likely coming from Western Africa (David and Capy 1988). It is doubtful that those introductions were unique events, and it is more likely that there were continuous events that followed recent human migration into the continent. Nevertheless, this differential pattern of introduction in the North and South is hypothesized to have created an ancestry cline along the East Coast (Kao et al. 2015; Bergland et al. 2016), hampering the identification of clines driven by spatially varying selection.

Despite those confounding factors, there is strong evidence that selection is acting to maintain at least some of the genotypic clines. Alleles of genes such as alcohol dehydrogenase (*Adh)* and glucose-6-phosphate dehydrogenase (*G6pd*) are clinal towards the same direction both in North America and Australia (Bubliy et al. 1999). This consistency between different continents suggests adaptation, likely driven by temperature. Temperature may represent a key factor in the existence of latitudinal clines, as many clinal phenotypes are also seasonal in temperate regions (Behrman et al. 2015). There is an agreement between seasonal and clinal alleles, with variants more common in the south of North America increasing in frequency just after the summer in temperate regions (Bergland et al. 2014; Cogni et al. 2014; Cogni et al. 2015; Rodrigues et al. 2021). Because of this relationship with temperature, it was proposed that shifts in certain clinal variants over time were related to global warming (Umina et al. 2005; Rodríguez-Trelles and Tarrío 2024). Nevertheless, there has not been a comprehensive investigation of changes in genome-wide clines over longer periods of time.

Historical samples of *D. melanogaster* have been analyzed in a few previous studies. Studies focusing on a single population (or a small group of geographically close samples) (Veeramah et al. 2020; Lange et al. 2022; Shpak et al. 2023) were valuable for elucidating demographic changes and identifying targets of selection. Other studies examined multiple historical samples scattered along the East Coast of North America (Cogni et al. 2014; Kapun et al. 2016; Cogni et al. 2017). However, these only targeted specific SNPs or genetic markers, which revealed different outcomes with different clinal patterns. For instance, clinal SNPs in the *cpo* gene were stable between 1997 and 2009/2010, while other clinal SNPs were lost or gained (Cogni et al. 2014; Cogni et al. 2017). Using historical genomic data that capture millions of polymorphisms could provide a more comprehensive understanding of the demographic and selective dynamics shaping clinal variation.

Looking at the whole genome of populations spread across the latitudinal gradient of the East Coast of North America with temporal samples could shed light on the processes at play in the establishment and maintenance of these clines. When tracking clinal changes over the long term, different patterns may occur: clines may become less steep, with populations becoming more genetically similar across the gradient; they may become steeper, with alleles associated with the north increasing in frequency in northern populations and alleles associated with the south increasing in southern ones; clines could reverse, with northern-associated alleles becoming southern-associated and vice versa; the entire cline may shift, with one allele rising in frequency across the whole gradient; or they may remain stable. Environmental changes between the sampled years, such as those driven by climate change, could favor shifts in clinal alleles. If global warming is driving adaptation, we expect to see an increase in the frequency of variants associated with warmer, lower latitudes. This effect would likely be most pronounced in SNPs within regions linked to genes involved in climate adaptation, which may constitute the largest proportion of clines impacted during this period (Rodrigues and Cogni 2021).

On the other hand, the observed pattern will also depend on the key processes that drove the formation and maintenance of the clines. If clines were generated by neutral processes, such as isolation by distance or secondary contact between two different genetic backgrounds, we would expect them to become less steep due to gene flow and subsequent mixing between populations, given that population sizes are high (Sprengelmeyer et al. 2020) and long-range dispersal is likely to be common (Coyne and Milstead 1987). Moreover, this cline weakening should be more prominent in neutral loci, while clines related to spatially varying selection would remain stable provided that selection pressures are consistent.

To study spatiotemporal genetic differentiation in *D. melanogaster*, we analyzed pool-seq genomic data from natural populations along the East Coast of North America. The samples were initially collected in 1997 and then recollected in 2009/2010, with a few additional collections in 2017 and 2022/2023. All samples were preserved frozen during the period that passed between collection and sequencing. The time span represents roughly 250 to 350 fly generations, assuming ten generations per year in the north and fifteen in the south (Pool 2015), where warmer conditions allow faster generation turnover. To put into perspective, this would be equivalent to 6,200 to 8,750 years in human generations. Thus, this is a powerful dataset to study processes shaping genetic variation along this latitudinal transect. Specifically, we asked whether spatial genomic differentiation and clinality changed over time, whether the persistence of clinal variation depended on genomic features such as recombination rate and chromosomal inversions, and whether loci showing persistent spatial differentiation or temporal signatures of selection were enriched for particular functional categories. In summary, we found that there was less spatial structure in 2009/2010 when compared to 1997, with most clines being lost over time. The clines that persisted were increasingly enriched at functionally relevant sites, consistent with a role of selection in maintaining clinal variation. Moreover, we identified signatures of selection in a few interesting genes (*e.g.*, insecticide resistance loci), which we unveiled using a window-F_ST_ approach.

## Results

### Genetic variation and population structure of fly populations across space and time

To understand how genetic variation has changed over time, we obtained pooled whole-genome sequencing data from 20 population samples along the East Coast of North America collected at four time points (1997, 2009/2010, 2017, and 2022/2023). Direct descendants of wild-collected flies were sampled and frozen until pooled sequencing (methods). Most samples were obtained in 1997 or 2009/2010 (Figure 1A, Table S1), mainly in late summer or fall. The exceptions consisted of three Florida populations sampled in December, two in 1997, and one in 2023. Because seasonal adaptation may influence our comparisons (Bergland et al. 2014; Behrman et al. 2015; Machado et al. 2021; Rodrigues et al. 2021), we added to our data set a Florida sample from December 2010 (Bergland et al. 2014; Machado et al. 2021). As a result, samples were seasonally aligned. Moreover, to increase space sampling coverage, we included two other populations from DrosRTEC, one from South Carolina and another from Georgia, both originally from 2010 (Bergland et al. 2014) (Bergland et al. 2014; Machado et al. 2021)(Bergland et al. 2014). Because there was slight variation in both sequencing depth and the number of flies per pool, we downsampled the sequencing data to achieve an average effective number of chromosomes (NE) of 20 per pool, ensuring comparability among samples (see Methods, Table S2, Figure S1). We used transversion to transition ratio (Ti/Tv), computed across all identified SNPs, as a measure of sample quality. Ti/Tv was on average 2.39 and similar across all samples (Table S3). The final dataset comprised 2,270,411 biallelic SNPs in the five major chromosomal arms (2L, 2R, 3L, 3R, X).

**Figure 1:**
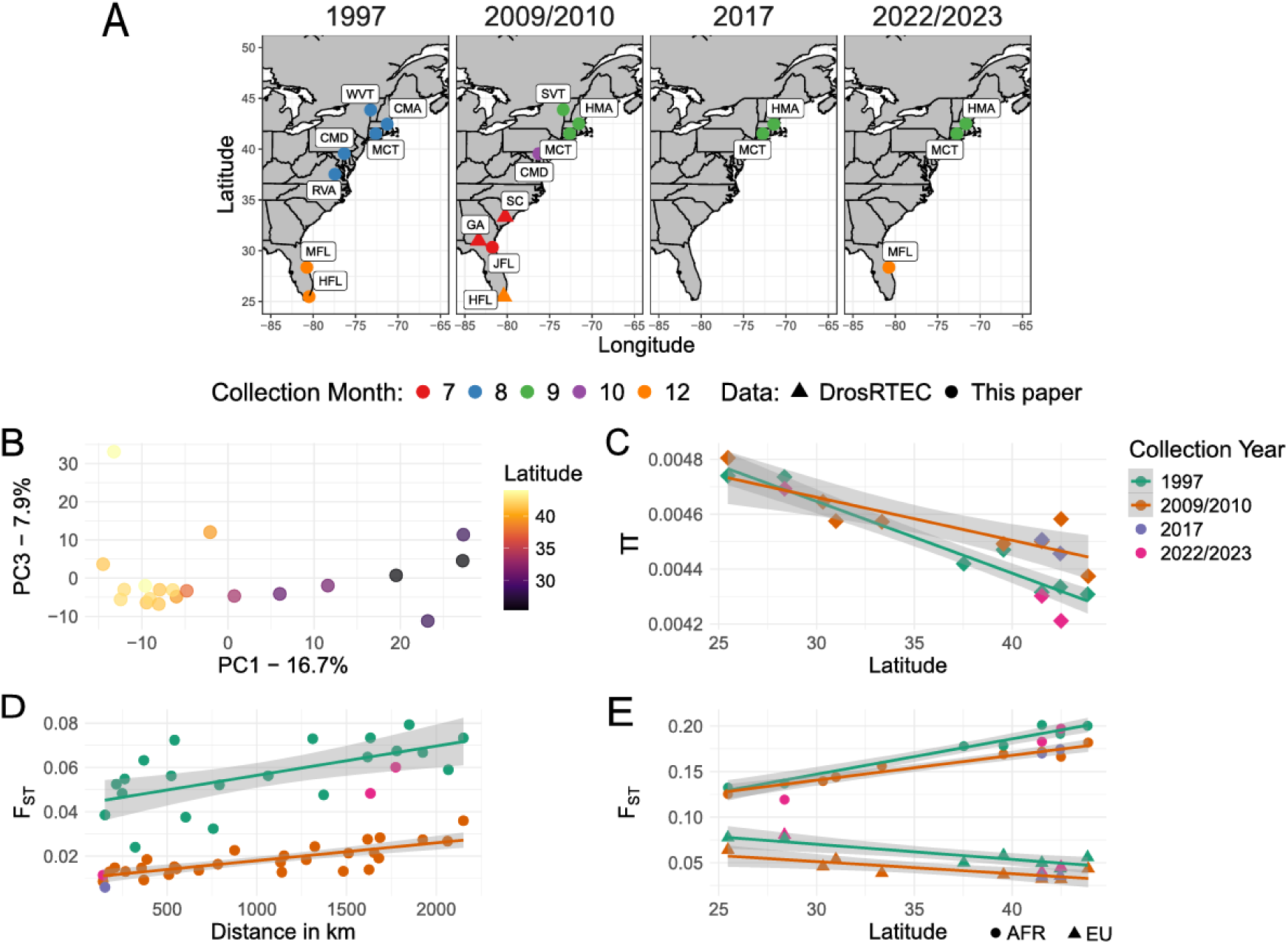
A - Populations sampled over time and space. Samples covered more than 25 years, which, assuming ten generations in the north and fifteen in the south, represents 250 to 350 generations in the north and south, respectively. Populations were sampled for genome-wide pool-seq, all pools first published here (circles) were composed of two females from each isofemale line, whereas publicly available data (triangles) were composed of one male per isofemale line. Sample names stand for the name of the city, followed by the name of the state where the samples were collected, or just the name of the state. B - Principal Component Analysis showing PC 1 and 3. C - Nucleotide diversity (π) of each pool; each point represents a population, colors reflect the collection year according to legend in the graph. D - Isolation by distance of each year; each point represents a pairwise F_ST_ of autosomes between populations sampled within the same year; years are colored according to legend. E - F_ST_ between the US populations and the Western-Central Africa (circles) or the European populations (triangles) using autosomes. For C, D and E regression lines are drawn for 1997 and 2009/2010 populations, grey areas show 95% confidence interval.

We conducted a principal component analysis (PCA) using all samples, which suggested PC1 was correlated with latitude (Figure 1B). The same pattern was observed for each autosomal arm when we did a per-chromosome PCA and when we used only samples from 1997 or 2009/2010 (Figures S2 and S3). Thus, latitude seems to explain most of the variation contained in the data set. We ran a linear model of the first 4 PCs against different environmental and collection-related variables to understand what influences our samples’ variation (Table S4). PC1 is primarily explained by latitude (*F(1, 18) = 169.00, p < .001, adjusted R^2^ = .898*; latitude effect: *β = −2.04, se = 0.16, t = −13.0*), but is also explained by collection month (*F(1, 18) = 13.72, p = .0016, adjusted R^2^ = .40*; collection month effect: *β = 5.5, se = 1.48, t = 3.7*) and the minimum temperatures from the three months prior to collection (*F(1, 18) = 8.43, p = .0094, adjusted R^2^ = .28*; minimum temperature effect: *β = 2.47, se = 0.85, t = 2.9*). This is unsurprising, as shifts in allele frequencies associated with seasonal changes are well-documented and may reflect adaptive tracking (Bergland et al. 2014; Rudman et al. 2022; Bitter et al. 2024; Nunez, Lenhart, et al. 2024). PC2 and PC4 are not significantly explained by any of our variables (Table S4), PC2 also only segregates sample CMD97B from the others (Figure S4). Sample CMD97B has a lower coverage depth than all other samples (18X), even after downsampling, and this might have had some influence on the allele frequency estimates. PC3 was partially explained by collection year (*F(1, 18) = 13.6, p = .0017, adjusted R^2^ = .40*; collection year effect: *β = −0.68, se = 0.19, t = −3.7*). Despite the small association between PC1 and collection month, most of its variation is related to the spatial distribution of populations.

To see how genetic diversity changed through the years, we computed nucleotide diversity (π) globally and, given known chromosomal inversion dynamics in natural *D. melanogaster* populations, we also computed it for each chromosomal arm per population. The diversity found in 2009/2010 here is slightly smaller than described in other papers with similar time and collection points (Fabian et al. 2012), their average π was 0.0061 in Florida and 0.0056 in Pennsylvania and Maine, while we found 0.0051 and 0.0048 in Florida and Vermont, respectively. We attribute this small difference to sampling variation (our samples had lower depths) and slight differences in data processing. Since we normalized sequencing depth to normalize NE among samples, we have no reason to suspect any bias related to the different time groups.

As expected, diversity decays in regions closer to the centromere (Begun et al. 2007; Langley et al. 2012), with a consistent pattern observed across the analyzed years (Figure S5). Using a linear mixed-effects model with chromosome as a random intercept to analyze all samples from all years, we found no significant tendency for changes in diversity across years (*β = 1.53E-06, s.e. = 1.71e-06, t = 0.894, p = .374*, Table S7, Figure 1C). Interestingly, however, when we exclude the years with fewer samples and analyze only populations from 1997 and 2009/2010, the year effect becomes significant, with populations from 2009/2010 being more diverse than populations from 1997 (*β = 9.62e-05, s.e. = 3.37e-05, t = 2.857, p = .006*, Table S7). Diversity was also negatively correlated with latitude (in both models), with samples from the north being less diverse (*β = −2.07e-05, s.e. = 2.43e-06, t = −8.549, p < .001*). We obtained similar results when we removed sample CMD97B from the analysis (Table S7). We repeated the analysis with each chromosomal arm separately, and the general pattern of decreased diversity in higher latitudes was similar across all of them, and no difference between years was found for any of them (Table S8, figure S6).

To better understand how the pattern of genetic differentiation changed over the years, we computed pairwise genome-wide F_ST_ (Figure S7). For between-year comparisons, we considered only years with sufficient sample sizes (at least four populations) to reduce outlier effects and ensure robust model estimates; thus, analyses were restricted to samples from 1997 and 2009/2010. On average, F_ST_ values on the autosomes were 0.039 smaller in 2009/2010 than in 1997 (multiple linear regression of distance and collection year, overall model *F(2,46) = 136, adjusted R^2^ = .849, p < .001*, year effect: *s.e. = 0.00247, t = −15.617, p = < .001*, Table S9). We also looked at the patterns of isolation by distance at both time points. For the autosomes, the relationship between F_ST_ and distance did not vary with collection year (there is no interaction with collection year, *F(1, 45) = 1.90, p = .175*), there was just an overall reduction in F_ST_ values between samples from 2009/2010 (Figure 1D), with both time periods presenting isolation by distance patterns (*β = 1.05e-05, s.e. = 1.901e-06, t = 5.517, p = < .001*). This holds even when excluding the CMD97B sample (the sample that was segregated in PC2, Table S9), though it seems that this removal increases the rate of F_ST_ by distance in 1997. We repeated the analysis for each chromosomal arm individually, including the X chromosome, which resulted in similar patterns for all of them (Table S10). The F_ST_ results from 2009/2010 agree with the F_ST_ of other populations collected in similar locations and times (Fabian et al. 2012). A previous study surveyed populations in 2000 and found similar F_ST_ values to the ones reported here for 1997 (Reinhardt et al. 2014). Thus, it seems clear from the data that the mean F_ST_ of 1997 is considerably greater than the mean of 2009/2010, which indicates that the populations have become more similar in recent years.

To understand if ancestry patterns changed over the years, we used available sequence data from populations of Central Western Africa and Europe as points of comparison to our samples (Table S11, see methods). We estimated the F_ST_ of the autosomes between our samples and each reference population (Figure 1E). As expected, in both of our most sampled periods (1997 and 2009/2010), F_ST_ with the African population increased as latitude increased. This increase rate was greater with samples from 1997 (0.004 F_ST_ units for each degree in latitude compared to 0.003 for 2009/2010, following a multiple regression analysis for collection year and latitude, *F(3, 11) = 99.84,* adjusted *R^2^ = .97, p < .001*, Table S12). The opposite effect was observed for F_ST_ with the European population: as latitude increased, F_ST_ decreased (0.0015 decrease in F_ST_ units for each degree of latitude, multiple regression analysis for collection year and latitude (overall model: *F(2, 12) = 27.29, R^2^ = .79 p < 0.001*; latitude effect: *s.e. = 0.00026, t = −5.80, p < .001*, Table S11). While the rate of latitudinal decrease in F_ST_ with European populations is equal for both analyzed periods, there was a shift in 2009/2010 to values on average 0.017 F_ST_ units smaller (year effect: *s.e. = 0.00337, t = −5.02, p < .001*, Table S11). Despite this trend, our less sampled periods (2017 and 2022/2023) do not appear to follow it, with values showing no clear pattern. Nevertheless, at least for African F_ST_ autosomal comparisons, it seems that North populations from 2009/2010 were especially more similar to the African population when compared to North populations from 1997. We repeated the analysis with each chromosomal arm separately. The general pattern was the same across all chromosomes (Figure S8A, Table S13), with two exceptions: on chromosome 3R, there were no significant differences between the F_ST_ values with the African population between the years (multiple regression analysis for collection year and latitude, overall model: *F(2,12) = 26.26, p < .001, adjusted R^2^ = .78*; year effect: *β = −0.013, t(12) = −1.42, p = .181*, Table S12), and on chromosome X, there was no relationship between F_ST_ with the European population and latitude (multiple regression analysis for collection year and latitude, overall model: *F(2,12) = 5.544, p = .02, adjusted R^2^ = .394*; latitude effect: *β = −0.0003, t(12) = −0.549, p = .59*, Table S12). We also computed global ancestry with intercept-free linear models as described by Bergland et al. (2016), and the results were very similar regarding latitude, but there is no significant variation between years (for European ancestry: overall model *F(4,15) = 42.1, p < .001, adjusted R^2^ = .918*, year effect *β = 0.009, p = .419*; for African ancestry: overall model *F(4,15) = 41.7, p < .001, adjusted R^2^ = .918*, year effect *β = −0.002, p = .872*; see Figures S8B, S9, and Table S14). Additionally, we inferred the European admixture proportions using the f4 ratio, which also revealed the same tendency as the intercept-free linear model did (Figure S8C). Altogether, this corroborates the existence of an ancestry cline within the East Coast of North America.

### Fewer Clinal Variants Observed in 2009/2010

To assess the presence of clines at SNPs, we used a general linear model of the frequency of each SNP against the latitude weighted by the effective number of chromosomes. We fitted those models on populations from 1997 and 2009/2010 separately. We examined 1,543,331 SNPs that passed the quality thresholds (see methods), both in 1997 and 2009/2010. The distribution of p-values shows an excess of small p-values in 1997 compared to 2009/2010 (Figure S10). With the p-values of those tests, we computed the number of clinal SNPs under a False Discovery Rate (FDR) that varied from 1% to 10% (Table S15, Figure S11). For an FDR of 10%, we found 277,632 clinal SNPs in 1997 and 55,766 in 2009/2010.

To determine whether this difference could be explained by unequal statistical power between the two sampling periods, we simulated clinal SNPs using each SNP’s depth and number of flies of each pool-sequencing. Slope values varied from 0 to 1. Applying the same FDR approach to those simulated datasets revealed a slightly higher power to detect clinal SNPs in 2009/2010 than in 1997 (Figure S12). Because NE was controlled between periods, we hypothesized that this difference might instead reflect the unequal number of populations sampled in each period. To test this, we repeated the analysis using matched datasets containing five populations from each period while controlling, as closely as possible, for geographic location and collection month. Specifically, we kept only populations from Vermont, Massachusetts, Connecticut, Maryland and Homestead, Florida. When we matched the populations in each period, the difference in power between the periods disappeared (Figure S12). We also looked at the false positive rate in those simulations, which were very low in all of them (mean of 0.014 using matched data for both periods, Figure S13).

Because of this difference in power, all the forward analysis of clinal SNPs focused on the matching samples dataset. We analysed 1,248,924 SNPs in this dataset, and still obtained more clinal SNPs in 1997 than in 2009/2010 (108,555 in 1997 and 20,689 in 2009/2010 at 0.1 FDR, Figure 2A and B). To statistically test this difference across years, we used two approaches: first we did a KS-test comparing the p-values distribution of both periods, which was significant (asymptotic two-sample Kolmogorov-Smirnov test, D^+^ = 0.072719, p < .0001); second, we performed a McNemar’s Chi-square test comparing the stability of clinal SNPs both at 10% and 5% FDR (McNemar’s chi-squared = 87864, df = 1, p < .0001 and McNemar’s chi-squared = 51905, df = 1, p-value < .0001, respectively). Only about 42% of clinal SNPs in 2009/2010 were clinal in 1997 (Table S16), indicating the emergence of new clinal SNPs. Although new clinal SNPs emerged, these gains did not compensate for the loss of previously clinal SNPs, resulting in an overall decrease in the number of clinal SNPs over the analysed period.

**Figure 2:**
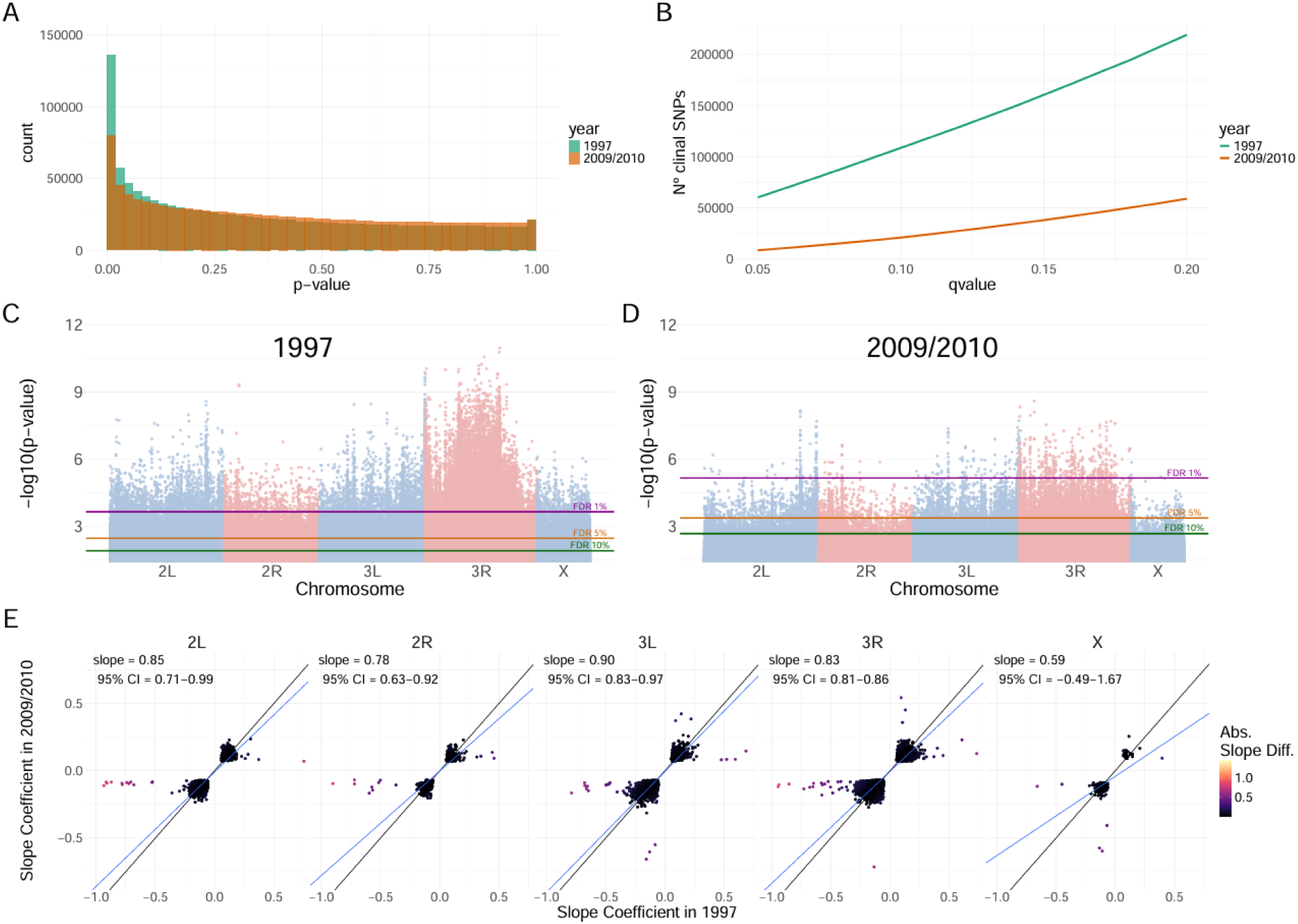
Number and steepness of clinal SNPs decreased in 2009/2010 when compared to 1997. A - Distribution of P-values based on a binomial GLM weighted for the number of flies in the pool and coverage depth, colored by period; the distribution of p-values of 1997 is skewed toward smaller values when compared to 2009/2010. B - number of SNPs considered clinal using different q-values cutoffs in each year. C (1997) and D (2009/2010) show the distribution of P-values across the 5 major chromosomal arms. E - Deming regressions of slope coefficients in 2009/2010 and 1997 of SNPs that were called clinal at both periods (at 15% FDR).

To check if this pattern of fewer clinal SNPs in 2009/2010 was independent of chromosomal arm, we computed the number of clinal SNPs under different FDRs for each chromosomal arm independently (Figure S14). The general pattern of more clinal SNPs in 1997 was still found in all chromosomes, but this pattern was much stronger in chromosome arm 3R. We then looked at the distribution of those p-values along the genome. While low p-values seemed uniformly distributed in 2009/2010, we found numerous low p-values in chromosomal arm 3R in 1997 (Figure 2C-D), likely related to inversion dynamics in this chromosome.

Focusing on SNPs that were clinal in both periods, we found that clinal slope concordance (alternative allele frequency increasing or decreasing in the same direction) was high (98.2% with a q-value of 0.2 to 99.9% with a q-value of 0.05 and similar to the all-samples dataset, Table S16). Using those concordant SNPs, we examined clines that shifted, that is, clines that had one of the alleles with higher or lower frequency along the entire latitudinal gradient. Those shifts could have occurred in two different directions: a decrease in the southern variant, which we will call a ‘North’ shift (because it is the northern variant that is increasing); or an increase in the southern variant, which we will call a ‘South’ shift. We wanted to understand if there was a bias toward ‘North’ or ‘South‘ types of shifts. If climate change–related warming were playing a role in the clinal changes over time, it would be reasonable to expect more ‘South’-type shifts than ‘North’-type shifts. To investigate this, we selected the top 50% of SNPs with the smallest differences in clinal slopes and refitted the GLMs, this time including samples from both tested periods (1997 or 2009/2010) in the same models and using it as a categorical predictor. Applying a 10% FDR to the year-related p-values identified 456 SNPs, of which 400 showed North shifts and 56 showed South shifts. However, examining the distribution of these shifts across chromosomal arms revealed that this pattern was particularly pronounced on chromosome 3R, where nearly 14 times more North shifts than South shifts were observed. Chromosomes 2L and 3L also showed approximately twice as many North shifts as South shifts, whereas chromosome 2R showed nearly equal numbers of shifts in either direction (Table S17). These chromosome-specific patterns may be influenced by shifts in chromosomal inversion frequencies, as discussed in the next section.

Focusing on the concordant SNPs, we examined the pattern of slope changes over time. To understand this, we conducted a deming regression analysis of the slope of clinal SNPs in 1997 and in 2009/2010. The relationship between slope coefficients from 2009/2010 and 1997 deviated significantly from a 1:1 relationship, with the deming regression slope of 0.865 (95% CI: 0.835- 0.896; H0:β=1, p < .001, Figure S15A). The increase rate was smaller than 1, indicating that, in general, slope values in 1997 were higher than in 2009/2010, suggesting a possible weakening of the clines over time. To see if this pattern was present regardless of chromosomal arm, we rerun the deming regression of clinal slopes for each chromosomal arm independently. After filtering for concordant SNPs using a 10% FDR, we could not reject the null hypothesis of a 1:1 relationship between slopes from the two years for chromosomes 2 and X (Figure S15B). In contrast, chromosome 3 showed evidence of changes consistent with a weakening of clines, particularly on the 3R arm, potentially reflecting changes in the frequencies of chromosomal inversions. Because these analyses were performed separately for each chromosome and some chromosomes contained relatively few SNPs identified as clinal in both periods at a 10% FDR (260 concordant SNPs in 2R and 1,009 in 2L), we repeated the Deming regressions using a slightly less stringent FDR threshold of 15%, which increased the number of concordant SNPs to 633 in 2R and 2,141 in 2L. At this threshold, the relationship between clinal slopes was consistently lower than 1:1 across all autosomes (Figure 2E), suggesting a general weakening of clines over time.

### Most Chromosomal Inversions Changed Over Time

Chromosomal inversions are common in *D. melanogaster*. Because they can suppress recombination in heterozygotes, it has been proposed that they play a major role in local adaptation (Kirkpatrick and Barton 2006; Kapun et al. 2016). They can also influence genetic diversity on the entire chromosomal arm (Corbett-Detig and Hartl 2012). Many of these inversions show clinal patterns that are convergent across continents (Kapun and Flatt 2019) or linked to other gradients, such as altitude (Pool et al. 2017). To investigate changes in their frequencies over time, we computed inversion frequencies using SNP markers (Kapun et al. 2014) for the seven most prevalent *D. melanogaster* inversions (Figure 3, Table S18). Our estimates for 1997 align with those made by Sezgin et al. (2004), who used the same collections but estimated inversions through PCR of their breakpoints. Our 2009/2010 estimates also match those reported for a similar time and location by Kapun et al. (2016).

**Figure 3:**
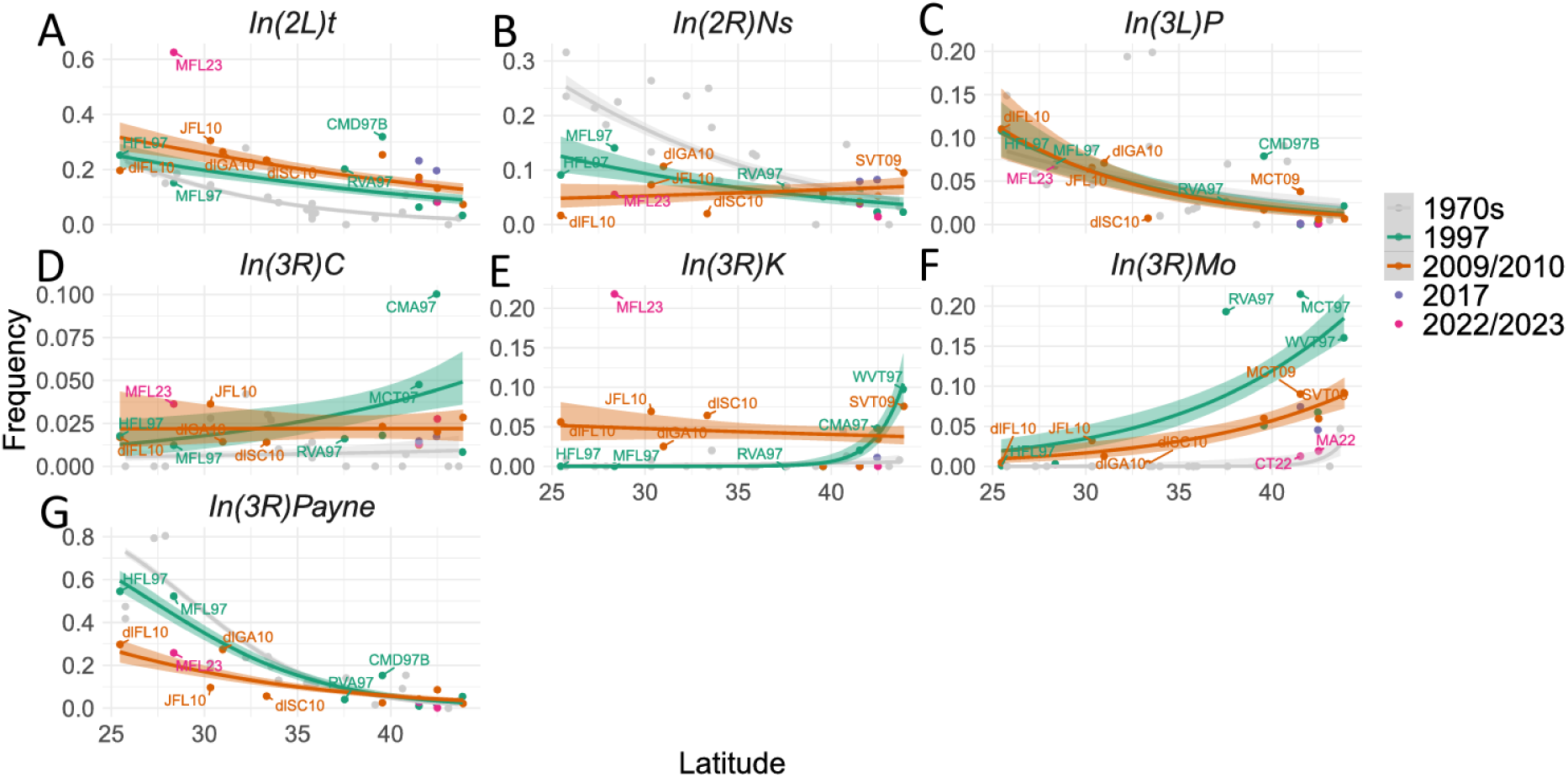
Inversion frequencies of the 7 major cosmopolitan inversions of *D. melanogaster*. 1970s data was retrieved from Mettler et al. (1977). Lines show the binomial glm with a logit link function, drawn only for samples collected in 1997 or in 2009/2010 as later collections lack sufficient spatial coverage for clinal estimates, grey areas show a confidence interval of 90%. For each model, coefficients (β), z-statistics, and *P*-values are reported for latitude and collection period, respectively: In(2L)t, β = −0.0639 (z = −8.10, *P* < .001) and β = 0.373 (z = 3.42, *P* < .001); In(2R)Ns, β = −0.0315 (z = −2.66, *P* = .008) and β = 0.00905 (z = 0.056, *P* = .955); In(3L)P, β = −0.126 (z = −7.75, *P* < .001) and β = −0.0303 (z = −0.139, *P* = .890); In(3R)K, β = 0.0549 (z = 2.75, *P* = .006) and β = 0.399 (z = 1.82, *P* = .068); In(3R)Payne, β = −0.181 (z = −18.26, *P* < .001) and β = −0.708 (z = −5.35, *P* < .001); In(3R)Mo, β = 0.131 (z = 7.38, *P* < .001) and β = −0.816 (z = −5.45, *P* < .001); and In(3R)C, β = 0.0425 (z = 2.02, *P* = .043) and β = −0.473 (z = −1.96, *P* = .050).

Three inversions remained somewhat rare and changed haphazardly: *In(2R)Ns*, *In(3R)C*, and *In(3R)K* (Figures 3B, 3D and 3E). *In(2R)Ns* changed direction completely: whereas it was more common in the South in 1997, it became more common in the North in 2009/2010. *In(3R)C* was practically absent in the South and rare in the North in 1997; in 2009/2010, it increased in frequency in the South and decreased in the North, but overall kept its frequency low. Something similar happened to *In(3R)K*, it was not present in the South and had small frequencies in the North in 1997, but in 2009/2010, it kept its frequency low along the latitudinal gradient. Interestingly, however, it reached a frequency of 22% in Florida in 2022.

*In(3L)P* is the most stable of inversions, and indeed its frequency pattern along the latitude did not change over the years (Figure 3C). Notably, when Kapun et al. (2016) used data from 1970 (retrieved from (Mettler et al. 1977)) and 1997 (retrieved from (Sezgin et al. 2004)) and compared them to their 2009/2010 samples, they saw a change in *In(3L)P* driven mainly by the 1970s data, with frequencies from 2000 and 2009 practically identical, compatible with our estimates.

For *In(2L)t*, we found an increase in the intercept between 1997 and 2009/2010 (Figure 3A), consistent with the consecutive increases found by Kapun et al. (2016) since 1970. However, we saw no change in the 2017 and 2022/2023 Northern samples. Though in the South there was a striking rise in its frequency, which went from close to 30% in 2009/2010 to over 60% in the 2023 MFL23 (Merritt Island, Florida, in 2023) sample.

Although *In(3R)Mo* frequency was consistently low in the South, it strongly decreased in the North in recent years (Figure 3F). This seems to be an ongoing pattern, as its frequency in 2017 and 2022 also decreased. It is likely that *In(3R)Mo* emerged after the out-of-Africa migration of *D. melanogaster* and has a cosmopolitan origin (Corbett-Detig and Hartl 2012), which might help explain its higher frequencies in the north. It was practically absent in Raleigh, North Carolina in the 1970s (Mettler et al. 1977), but increased to 18% by 2003 (Langley et al. 2012). This is not necessarily in conflict with what we found, as our time frames do not align perfectly.

In contrast to *In(3R)Mo*, *In(3R)Payne* remained stable in the North but decreased its frequency in the South (Figure 3G). Kapun et al. (2016) also examined this inversion, their frequency estimates from 2000 and 2010 match ours from 1997 and 2009/2010. However, they found no temporal changes in the slope or intercept of this inversion. Notably, the increase in the intercept of this inversion cline between 1979 and 2004 in Australia (Anderson et al. 2005; Umina et al. 2005) does not match what we or Kapun et al. (2016) observed.

### Recombination narrows clinal variation, leaving functionally enriched SNPs

To determine whether clinal SNPs were disproportionately represented in specific genomic regions and whether this pattern changed over time, we calculated the odds ratio comparing the odds of a clinal SNP occurring in a given genomic region with the corresponding odds for 100,000 randomly sampled non-clinal SNPs. We repeated this random sampling 10,000 times, recalculating the odds ratio for each random sample to generate a distribution of odds ratios expected under random sampling (Figure 5). We considered a genomic class enriched or depleted when its odds ratio distribution was consistently above or below the expected value of 1.

**Figure 5:**
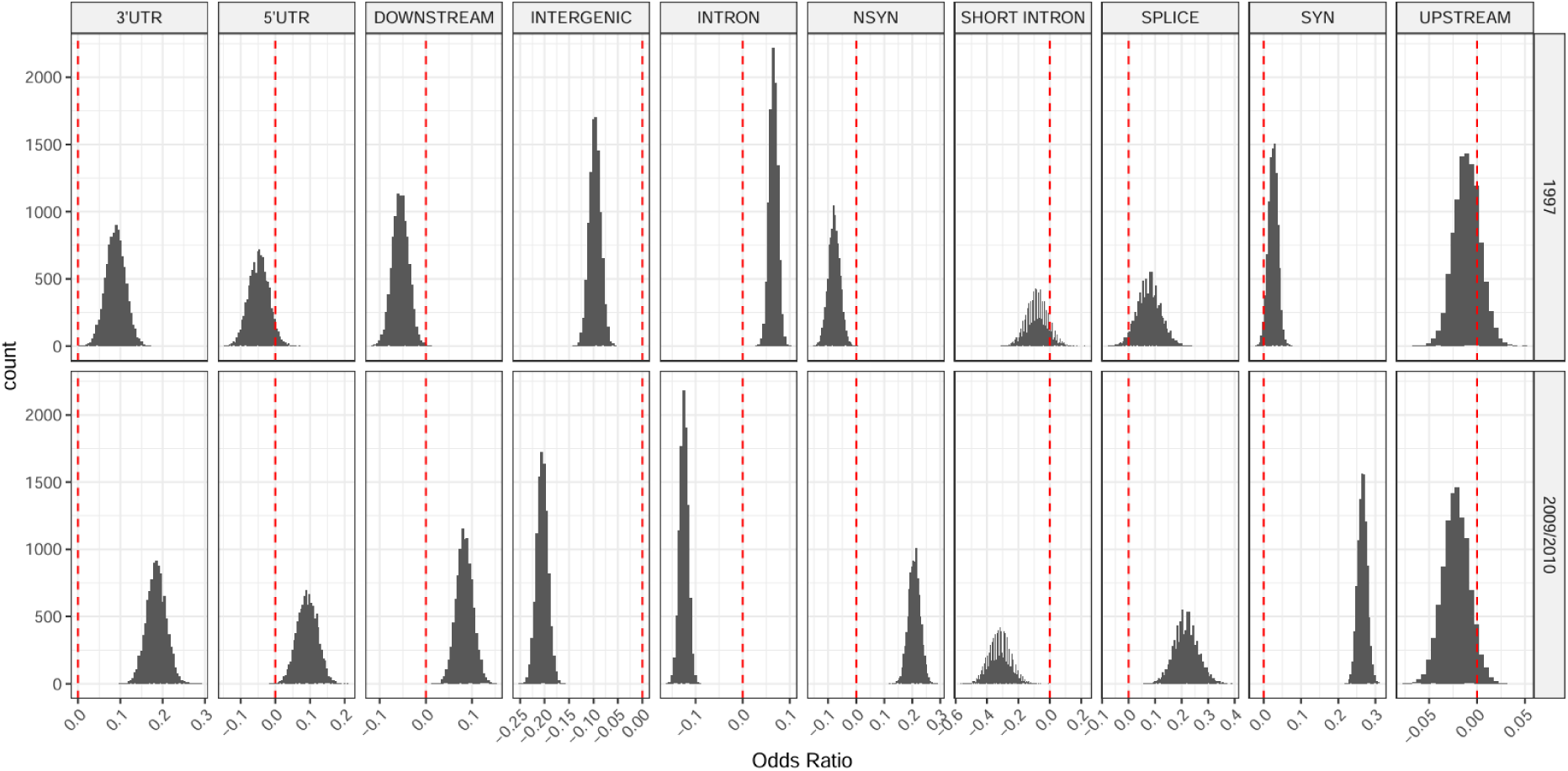
Odds ratio of a clinal SNP being in a specific region vs a set of 100.000 random drawn non-clinal SNP being in the same region. The random set of non-clinal SNPs was resampled 10.000 times (single grey bars). Red dashed line shows equal chances of clinal vs non-clinal SNP in that genomic region. We used a q-value of 0.1 to consider a SNP clinal.

In 1997, using a 10% FDR cutoff, intergenic and non-synonymous SNPs were depleted among clinal SNPs, whereas intronic SNPs were enriched. Synonymous SNPs, SNPs in 3′ UTRs, and SNPs in splice-related regions also showed a tendency toward enrichment, while downstream regions showed a tendency toward depletion. In 2009/2010, intergenic, intronic, and short-intron SNPs were depleted among clinal SNPs, whereas 3′ UTR, downstream, non-synonymous, synonymous, and splice-related SNPs were enriched. 5′ UTR SNPs also showed a tendency toward enrichment. Using a 5% FDR cutoff to define clinal SNPs produced the same overall pattern (Figure S16C). Similarly, analyses using the entire dataset rather than the matched dataset, which included matched populations, yielded comparable results (Figures S16A and S16B).

We assessed the overlap between the distributions of log2 odds ratios across timepoints to determine whether the enrichment of clinal SNPs changed over time (Table S19). Intronic regions shifted from enrichment in 1997 to depletion in 2009/2010 (log2 OR = 0.066 vs. −0.124), with no overlap between the distributions. Intergenic regions remained depleted but showed stronger depletion in 2009/2010 (log2 OR = −0.097 vs. −0.204), also with no overlap. Short introns showed a similar shift toward stronger depletion (log2 OR = −0.080 vs. −0.317), although the distributions showed some overlap (8.1%). In contrast, non-synonymous regions shifted from depletion in 1997 to enrichment in 2009/2010 (log2 OR = −0.078 vs. 0.208), with no overlap between the distributions. Downstream and synonymous regions also shifted toward enrichment in 2009/2010 (downstream: log2 OR = −0.055 vs. 0.084; synonymous: log2 OR = 0.027 vs. 0.266), with no overlap. UTR and splice-related regions showed higher enrichment in 2009/2010, although their distributions overlapped to some extent (3′ UTR: log2 OR = 0.089 vs. 0.185, overlap = 3.4%; 5′ UTR: log2 OR = −0.046 vs. 0.093, overlap = 1.7%; splice-related: log2 OR = 0.078 vs. 0.217, overlap = 11.6%). Finally, 5′ UTR regions also shifted from slight depletion in 1997 to enrichment in 2009/2010. Overall, with the exception of downstream regions, clinal SNPs in more recent years showed increased enrichment in coding regions and splice-related sites, accompanied by greater depletion in intergenic, intronic, and short-intron regions.

We suspected that the reduction in the number of clinal SNPs between 1997 and 2009/2010 could be caused by gene flow along the latitudinal gradient. This gene flow would mainly drive the loss of neutral clinal SNPs (SNPs that were linked to other SNPs targeted by spatially varying selection). If this were the case, it would be reasonable to expect that most clinal SNPs that were lost between 1997 and 2009/2010 to be in regions of higher recombination. It would also be expected that genomic regions with lower recombination rates to be more densely populated by clinal SNPs than regions with higher recombination rates. Those expectations are based on the species’ natural biogeographic history in North America, where an ancestry cline exists, and therefore, most clines may not have initially arisen through natural selection. Conversely, if clines were initially the product of natural selection, clinal SNPs within regions of higher recombination rates would probably better resist the influence of gene flow than those within regions of lower recombination rates, where selection is less effective, and they would also be more frequent in regions of higher recombination rates.

To test whether regions with higher recombination rates lost more clinal SNPs between 1997 and 2009/2010, we identified SNPs that were clinal in 1997 and modelled their persistence as clinal in 2009/2010 as a function of the recombination rate of the genomic region in which they were located, including the genomic window used to calculate recombination rate as a random effect. Recombination rate had a small but significant positive effect on the probability that a SNP ceased to be clinal, with each one-unit increase in recombination rate increasing the odds of losing clinal status by approximately 25% (β = 0.227, odds ratio = 1.25, *p* < .001; Table S20, Figure S17A). We then repeated the analysis separately for each chromosome. The positive association between recombination rate and loss of clinal status was significant on 2L (β = 0.261, odds ratio = 1.30, *p* < .001), 2R (β = 0.162, odds ratio = 1.18, *p* = .018), and 3L (β = 0.321, odds ratio = 1.38, *p* < .001), but not on 3R (β = 0.029, odds ratio = 1.03, *p* = .277) and X (β = 0.305, odds ratio = 1.36, *p* = 0.124; Table S20, Figure S17B).

We then repeated the same model strategy using all SNPs to test whether clinal SNPs were more frequent in regions with lower recombination rates. For both periods, we found a significant negative relationship between recombination rate and the probability that a SNP was clinal (Table S21). In 1997, each one-unit increase in recombination rate was associated with a 0.94-fold change in the odds of a SNP being clinal (β = −0.060, p < .001; Figure S18A). When the analysis was repeated separately for each chromosome, the negative association was significant on chromosome 3L (OR = 0.85, p < .001) and chromosome X (OR = 0.96, p = .001), whereas no significant association was detected on 2L or 2R. Chromosome 3R showed a weak positive association between recombination rate and the probability of being clinal, although this effect did not reach statistical significance (OR = 1.04, p = .058; Figure S18B). In 2009/2010, the negative association was substantially stronger, with each one-unit increase in recombination rate corresponding to a 0.83-fold change in the odds of a SNP being clinal (β = −0.191, p < .001; Figure S18C). At the chromosome level, this negative relationship was significant on 2L (OR = 0.82, p < 0.001), 2R (OR = 0.92, p = .014), 3L (OR = 0.77, p < .001), and X (OR = 0.89, p = .001), but not on 3R (OR = 0.97, p = .303; Figure S18D). Altogether, these results indicate that clinal SNPs are more frequently found in regions of low recombination, and that this association became stronger and more widespread across chromosomes in 2009/2010.

### Spatially consistent selection targets detoxification genes across decades

To investigate the action of natural selection over time, we searched for signatures of selection on genomic variation at each location across different time points. The signature of selection we adopted is an excess of genetic differentiation, measured by 100 SNP windows F_ST_ and identified as the genomic regions with the highest F_ST_ values (top 0.1%). This approach assumes that the overall genomewide distribution of F_ST_ reflects neutral evolution, and the outlier windows, which deviate from the genomewide pattern, were more likely to have been targeted by selection. Then, we mapped the genes that overlapped within each of those outlier F_ST_ windows (Table S22). Next, searching for spatially consistent selection, we focused on regions that were outliers across locations for each pair of time points analyzed (i.e., 1997-2009/2010 and 1997-2022/2023).

In total, 14 genomic regions encompassing 27 genes were identified with some degree of overlap between outlier windows during the 1997 to 2009/2010 timeframe (Figure S19, Table S23). Still, no gene was present in the temporal F_ST_ outliers of more than two populations. Between 1997 and 2022/2023, four peaks, containing 20 genes (Table S23), from different geographical locations overlapped, with only one of those peaks present in all three pairs of populations (Figure 6). Across the two periods, 10 genes were shared, all of which were located within the same genomic region on chromosome 2R containing the *Cyp6a* gene cluster and *Cyp317a1*. This region was identified as a shared outlier in Connecticut and Florida between 1997 and 2009/2010, and in Connecticut, Massachusetts, and Florida between 1997 and 2022/2023, indicating that differentiation at this region has persisted across sampling periods and across multiple geographical locations. The shared region encompasses *Cyp6a22, Cyp6a17, Cyp6a19, Cyp6a20, Cyp6a8, Cyp6a9, Cyp6a21, Cyp6a23, Cyp317a1*, and *Pcf11*. Cytochrome P450 genes are involved in the metabolism of endogenous compounds and xenobiotics, including insecticides (Scott et al. 1998; Lu et al. 2021), and several genes in this cluster have previously been associated with insecticide resistance (Daborn et al. 2007)(Battlay et al. 2018; Duneau et al. 2018)(Daborn et al. 2007). In particular, disruption of *Cyp6a17* increases susceptibility to pyrethroids, the second most common class of insecticides (Battlay et al. 2018; Duneau et al. 2018). Three chimeric variants involving *Cyp6a17* and *Cyp6a23* have also been described, including two fusions of *Cyp6a17* with *Cyp6a23* and a third variant composed primarily of *Cyp6a23* with a small portion of *Cyp6a17* within the first exon (Good et al. 2014; Chen et al. 2024). Selection was inferred to favor the intact variant between 1975 and 1983 in Providence, Rhode Island (Lange et al. 2022).

**Figure 6:**
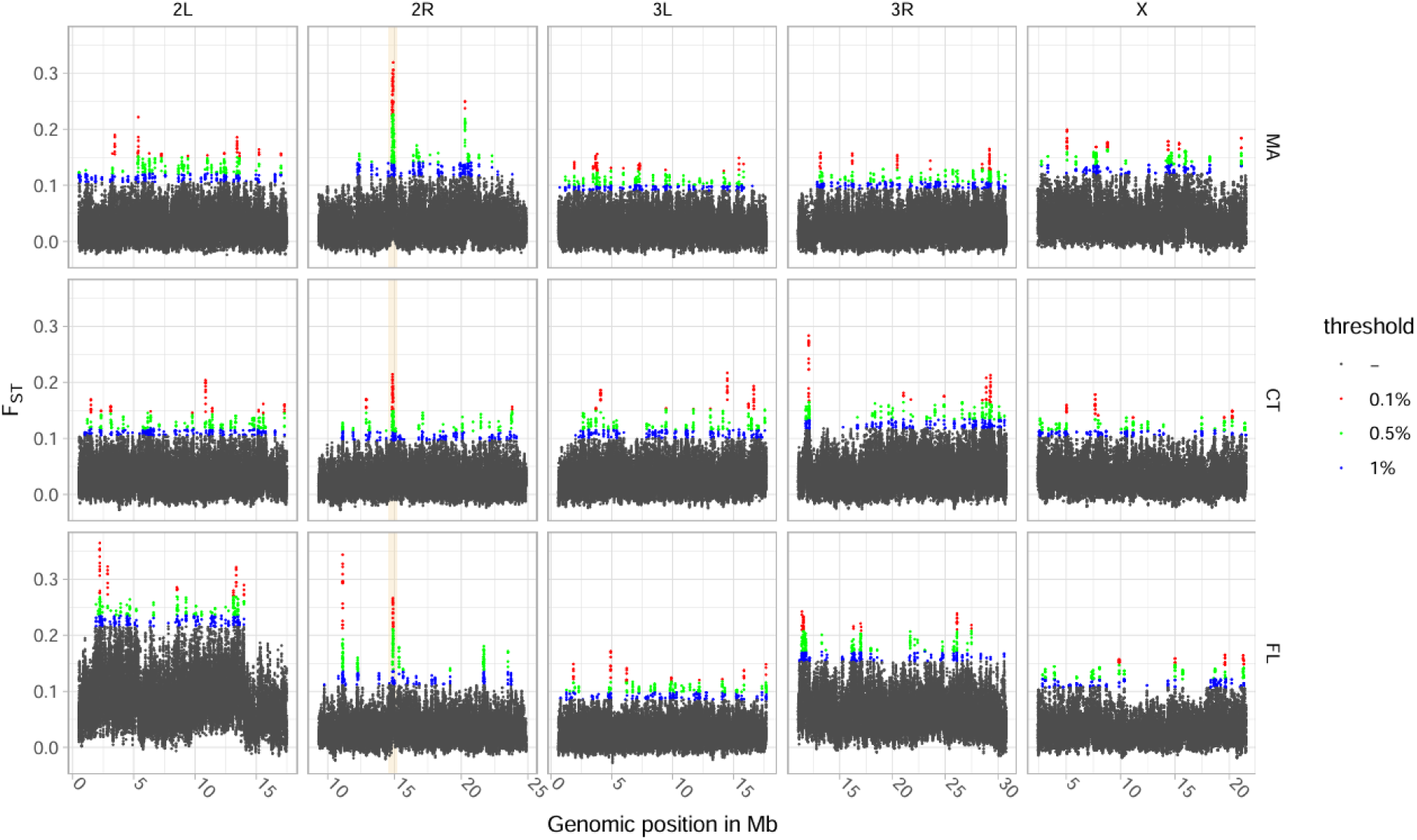
Window-F_ST_ scan between 1997 and 2022/2023. Windows were defined as 100 SNPs in length with a 10-SNP stride. Populations used were: Concord 1997 and Harvard 2022 for Massachusetts (MA); Middlefield 1997 and 2022 for Connecticut (CT); and Merritt Island 1997 and 2023 for Florida (FL). Highlighted area shows the only shared peak between all three pairs of populations and contains genes of the P450 gene family.

We wanted to investigate if selection was also favoring the intact *Cyp6a17* variant in our analyzed timeframe. Although it is difficult to infer structural variants from pool-seq data, we looked at the read depth of this genomic region to identify the possible presence of structural variants, as gene copy number or the chimeric variants described before (Figure S20). The chimeric variant predominantly lacking *Cyp6a17* appears to have been present in 1997 in Massachusetts and Florida (Merritt-Island), but this is less clear for Florida (Homestead) and Connecticut. By 2009/2010, the intact variant seemed prevalent in Massachusetts and Florida (Homestead) populations, while Connecticut exhibited the *Cyp6a17* deletion. Intriguingly, in 2022/2023, all populations analyzed (Massachusetts, Connecticut, and Florida - Merritt Island) showed doubled read depth in the *Cyp6a22* and *Cyp6a19* gene regions, which flank *Cyp6a17* and *Cyp6a23*. This suggests the emergence of an uncharacterized copy number variant that has increased in frequency in recent years. Regardless of this possible scenario, the intact *Cyp6a17* variant appears to be declining in frequency, contrary to trends observed between 1975 and 1983, potentially reflecting a shift in selection pressures acting on this locus.

Using the genes in the top 0.1% highest F_ST_ peaks, we performed a Gene Ontology (GO) analysis for each pair of populations. All terms with a p-value < 0.05 are available in Table S24. We then focused on enriched biological process terms shared among locations. Between 1997 and 2009/2010, 37 biological process terms were enriched in more than one location, whereas 13 were shared between locations when comparing 1997 with 2022/2023 (Table S25). These terms were not independent, as many shared genes with other enriched processes or represented closely related levels of the same biological pathway. For example, several of the terms enriched between 1997 and 2009/2010 described different aspects of vesicle docking and exocytosis and were driven by overlapping sets of genes. Despite this redundancy, several recurring functional categories were particularly notable. Processes related to detoxification were enriched across multiple locations in both periods. Specifically, ‘insecticide metabolic process’ and ‘toxin metabolic process’ were enriched in Massachusetts, Connecticut, and Florida between 1997 and 2009/2010, with *Cyp6t3* contributing to the enrichment in Massachusetts and *Cyp6a8* in Connecticut and Florida. In the comparison with 2022/2023, the same processes were again enriched in Connecticut and Florida, in both cases driven by *Cyp6a8*. Other processes associated with *Cyp6a8*, including ‘lauric acid metabolic process’, ‘medium-chain fatty acid metabolic process’, and ‘response to caffeine’, were also shared between locations in both periods. Another recurring signal involved transcription termination: both ‘DNA-dependent transcription, termination’ and ‘termination of RNA polymerase II transcription’ were enriched in multiple locations in both periods, consistently involving *Pcf11*, which is also located within the recurrent 2R outlier region. Together, these results indicate that several functional categories, particularly those related to detoxification and metabolism, were repeatedly associated with spatially shared F_ST_ outliers across sampling periods, although the specific genes contributing to these enrichments could differ among locations.

## Discussion

Latitudinal clines in *Drosophila melanogaster* are among the most studied of any species. Many lines of evidence suggest that clines are shaped by spatially varying natural selection, for instance, there is a parallelism in latitudinal clines across different continents (Reinhardt et al. 2014; Bergland et al. 2016) and in altitudinal clines (Klepsatel et al. 2014; Pool et al. 2017). Moreover, many *D. melanogaster* clines also show seasonal variation (Bergland et al. 2014; Rodrigues et al. 2021; Machado et al. 2021; Nunez, Lenhart, et al. 2024; Nunez et al. 2025), and *D. melanogaster* shares more clinal genes with its sister species, *D. simulans*, than expected by chance (Machado et al. 2016). However, in North America, the colonization pattern of *D. melanogaster* has led to an ancestry cline that hampers the identification of spatially selected variants. In this study, we used historical samples to revisit past clines and compare them to more recent ones. We found fewer clinal SNPs and more homogeneous populations in the recent samples, suggesting much of the past clinal variation was neutral and has since been erased due to gene flow.

Differentiation between populations decreased between 1997 and 2009/2010 (Figure 1D), as measured by F_ST_. This is consistent with a homogenization of populations due to gene flow. Moreover, there was an increase in nucleotide diversity when comparing populations from 2009/2010 to populations in 1997 (Figure 1C). We also observed a decrease in F_ST_ between the US populations and the presumed ancestral populations (Africa and Europe). Higher latitude populations experienced greater increase in nucleotide diversity and decrease in F_ST_ with Africa. This suggests there has been asymmetric migration along the US cline, with greater gene flow from South to North. On the other hand, ancestry clines, determined using both the linear model method (Bergland et al. 2016) and the f4-ratio, remained consistent throughout the years. Although this seems contradictory, it is possible that change in ancestry is too recent to affect both the f4-ratio and the linear model method (see Figure S2 in Bergland et al. 2016).

Although the pattern described so far suggests ongoing homogenization of populations, more recent samples yield unexpected results. First, we find that F_ST_ values between pairs of populations sampled in 2022/2023 are higher than those observed among populations from 2009/2010. This difference largely disappears when isolations by distance patterns are examined per chromosome, with the exception of arms 2L and 3R. We therefore suspect that the elevated differentiation is linked to the sharp increase in frequency of inversions In(2L)t and In(3R)K in the Florida 2023 sample (MFL23). Notably, In(2L)t has previously been associated with adaptive tracking of seasonal changes, and in particular with maximum temperatures before sampling (Nunez, Lenhart, et al. 2024). Because a Virginia population, described in Nunez et al (2024), does not show a comparable increase in these inversions, this may represent a population-specific shift rather than a general trend. Second, nucleotide diversity shows a different pattern. While diversity levels in 2017 are similar to those from 2009/2010, northern populations sampled in 2022/2023 exhibit reduced diversity, resembling values from 1997. If homogenization observed between 1997 and 2009/2010 had continued, we would expect diversity in 2022/2023 to be equal to or higher than that of 2009/2010. However, given that our dataset includes only three populations from this period, it remains uncertain whether this reduction reflects a broad latitudinal pattern or is restricted to the sampled populations. Another explanation could be variation in the strength of seasonal bottlenecks. We tested whether population differentiation was related to temperature in the months prior to collection and found no correlation. However, other variables not examined here may influence bottleneck strength (Machado et al. 2021; Shpak et al. 2023; Nunez et al. 2025).

Considering all chromosomal arms, chromosome X had the smallest proportion of African ancestry in the south. It is also the chromosome with the smallest correlation between geographic distance and F_ST_. This is somewhat unexpected, as the smaller effective population size of the X chromosome would theoretically increase drift and therefore increase F_ST_ (Wright 1931; Slatkin and Voelm 1991). We could also expect spatially varying selection to act more rapidly on this chromosome due to the faster-X effect. This would increase genetic differentiation, unless uniform selective pressure outweighs the spatially varying selection. A possible explanation for the smaller differentiation and overall lower African ancestry is that introgression of an African genetic background into a primarily European gene pool is harder on the X chromosome. This follows Haldane’s Rule, which says that genetic incompatibilities are more likely to appear in the heterogametic chromosome (Haldane 1922). Supporting this idea, the North Carolina DGRP population showed greatly reduced African ancestry in chromosome X when compared to the autosomes (Pool 2015), and reduced European introgression was found on the X among African flies (Pool et al. 2012). This initial barrier to the gene flow of the X chromosome, and its consequently weaker latitudinal ancestry gradient, could explain the lesser geographic differentiation of the X chromosome.

Our data showed a consistent reduction of clinal SNPs between 1997 and 2009/2010. Beyond the reduction in the number of clinal SNPs, there were very few changes in slope direction, with an overall decline in the clinal slope of SNPs that remained clinal when we disregarded the chromosomes. The prediction of slope reduction and loss of clines as time passes was proposed before (Bergland et al. 2016; Cogni et al. 2017; Rodrigues and Cogni 2021). This idea mainly anchored itself on the assumption that migration would be substantial enough to admix the flies from different populations along the latitudinal transect. This is especially plausible for *D. melanogaster* clines on the East Coast of North America because of its dual foundation process (Duchen et al. 2013; Kao et al. 2015; Bergland et al. 2016) and presumably high dispersion rates (Coyne and Milstead 1987). As a human commensal, *D. melanogaster* is also likely assisted by human transport. Moreover, its sister species, *D. simulans*, has a low capacity to overwinter (Boulétreau-Merle et al. 2003) and probably recolonize, at least partially, northern regions every year (Behrman et al. 2015; Machado et al. 2016). This high dispersion capacity of such a similar species indicates that *D. melanogaster* likely has a comparable ability. These results corroborate the idea that North American *D. melanogaster* populations are indeed connected and have exchanged migrants.

The reduction in clinal SNPs is not even across the genome. The results indicate that the inflation of clinal SNPs in 3R in 1997 and its absence in 2009/2010 are related to the inversion *In(3R)Payne*, and maybe to some degree to *In(3R)Mo*. *In(3R)Payne* decreased in frequency in 2009/2010 in the south, whereas it was strongly clinal in 1997. While *In(3R)Mo* was less frequent across the entire transect, it was also clinal in 1997 but decreased in frequency in 2009/2010 in the North. This indicates that clinal SNPs in 3R were probably linked to those inversions. Although the reduction in clinal variants was particularly pronounced in this chromosomal arm, it occurred across all chromosomes, indicating that the overall decline in clinal SNPs cannot be attributed solely to changes in inversions.

Many of the clinal SNPs observed in 2009/2010 were not clinal in 1997, indicating that they might represent newly emerging clinal SNPs. The adaptive effects of these new SNPs may have been previously masked by maladaptive neighboring variants and only became detectable after several generations of recombination. Perhaps this could be better explained by a more general pattern, as the different clinal enrichment pattern of those different years: there seems to be higher enrichment in regions of higher phenotypic impact in 2009/2010 than in 1997. This seems to be an indication that the remaining clinal SNPs are being maintained, and maybe reinforced, by selection, while the neutral clines tend to weaken or be lost.

Because clinal variants are often a product of natural selection, it is plausible that they would be more easily found in genomic regions with higher phenotypic impact. Still, many of those clines were probably not initially a consequence of spatially varying selection, but rather were present due to *D. melanogaster*’s dual colonization of North America. Over time, it would be expected that the influence of the initial colonization would lessen, and that the neutral regions would be less likely or be negatively enriched in more recent years. We found this pattern, with genomic regions traditionally regarded as neutral being less enriched for clinal SNPs in 2009/2010 than in 1997, and clinal sites within coding regions more enriched in 2009/2010 than in 1997.

Variants within coding regions, including both non-synonymous and synonymous SNPs, showed substantial changes in enrichment between periods. Non-synonymous SNPs were somewhat depleted among clinal variants in 1997 but became enriched in 2009/2010. Because non-synonymous SNPs can have some of the greatest impacts on phenotype, it is understandable that they may be subject to spatially varying selection, and why its clinality would be maintained by it. Synonymous SNPs only become enriched in 2009/2010 and, although there is a common assumption that they should be neutral because they do not impact the final structure of the protein, there is some evidence that they could also affect fitness. Strong negative selection was found acting on synonymous SNPs in *D. melanogaster* (Lawrie et al. 2013; Machado et al. 2020). A functional example of how synonymous mutations can affect the *D. melanogaster* phenotype is found in the *alcohol dehydrogenase* (*Adh*) gene, where shifts to less preferred codons have led to reduced *Adh* expression and decreased alcohol tolerance (Carlini and Stephan 2003; Carlini 2004). Many of those sites could be under spatially varying selection, which maintains their clines despite years of admixture. Alternatively, clinal synonymous sites may indirectly capture variation at other loci through linkage, such as non-synonymous or undetected structural variants that were not clinal in 1997 but became clinal by 2009/2010 and, owing to their lower frequencies, were not detected as clinal.

Splice-related regions also differed in their enrichment between periods, shifting from no significant enrichment in 1997 to enrichment in 2009/2010. Splice related regions, have also potentially strong phenotypic effects. Moreover, temperature has a substantial effect on splice parameters for many genes (Jakšić and Schlötterer 2016). An example of this is in the *period* (*per*) gene, which regulates day sleep in *D. melanogaster*. The splicing efficiency of a small intron in *per* 3’UTR (*dmpi8*) varies with temperature: cooler temperatures enhance splicing, reducing day sleep, while higher temperatures reduce splicing efficiency, increasing day sleep (Majercak et al. 1999). Notably, a haplotype linked to higher splicing efficiency is more common in temperate regions of Australia, whereas a haplotype linked to reduced splicing efficiency is more frequent in the tropical areas, promoting higher levels of day sleep in this population (Yang and Edery 2018). Clines affecting the splicing efficiency of this intron were also characterized in altitude gradients in Africa (Cao and Edery 2017). This example illustrates how changes in splicing can serve as targets of spatially varying selection. If this pattern is widespread across the genome, we would expect to see increasing enrichment of these regions among clinal SNPs over time, especially as non-targeted, neutral regions are mainly influenced by admixture in the absence of selective constraints.

Lastly, intergenic regions, introns were more depleted in 2009/2010 than in 1997. For many organisms these regions are regarded as somewhat more neutral, and having smaller phenotypic impact. However, there is compelling evidence that both intronic and intergenic regions are subject to purifying selection and adaptive evolution (Andolfatto 2005), probably due to the existence of numerous regulatory elements (Halligan and Keightley 2006). Short introns, however, are often used as neutral reference regions because they are the least constrained in *D. melanogaster* genome (Parsch et al. 2010; Clemente and Vogl 2012), and they were also more depleted in 2009/2010 than in 1997. Nevertheless, being subject to selection does not necessarily imply that these regions are frequent targets of spatially varying selection. If their functions are similarly constrained across different environmental contexts, they may be less likely to harbor variants whose fitness effects differ geographically, and therefore less likely to be maintained as clinal by spatially varying selection. If that is the case, such regions would be less likely to be maintained as clinal, which could account for their strong negative enrichment in 2009/2010.

In addition to different enrichment patterns of genomic regions, we observed that clinal SNPs in regions of higher recombination rate are more easily lost than in regions of lower recombination rate. Moreover, we saw that, independently of the analysed year, there were more clinal SNPs in regions of lower recombination rate. We recognise that the recombination rate estimates used in this study do not take into account the impact of chromosomal inversions. This is probably the reason why the effect of recombination is different between chromosomes. Nevertheless, those estimates should be suitable to detect at least the general relationship between recombination rate and maintenance of clinal SNPs. This general pattern suggests that as gene flow reshuffles the genome and linkage disequilibrium breaks down, the initially established clines in neutral regions are lost, leaving only those maintained by spatially varying selection.

Apart from spatially varying selection, we looked for signatures of selection focusing on spatially global targets using the available pairs of populations collected in the same locality but in different years. We found shared signatures of selection in regions related to xenobiotic resistance, especially in those of the P450 gene family. Copy-number variants in this gene family differed greatly between populations at opposite ends of North America and Australia (Schrider et al. 2016). Not only signatures of selection were found in another temporal study in North America (Lange et al. 2022), but it is also found when comparing different continents (Nunez, Coronado-Zamora, et al. 2024). This reinforces the idea that insects are suffering insecticide pressures around the world and that they tend to respond rapidly to it (Haddi et al. 2023). Moreover, from our examination of sequencing depth, it seems that different variants are responsible for this signal in different locations, which could point to different origins of possible resistance.

Undertaking a large-scale spatiotemporal analysis required that we handled several technical caveats to be able to better compare our samples. Our 1997 samples initially had less coverage depth than most of our more recent samples. We circumvented this problem by downsampling to a common effective number of chromosomes across samples. Whenever appropriate, we rerun our analysis without our sample with smaller depth coverage (CMD97B). Moreover, we used a heuristic caller (PoolSNP) that minimized single-sample errors of pooled data (Guirao-Rico and González 2021). Another consideration was the seasonal misalignment of some samples. We addressed this by adding or removing samples to ensure alignment on both season and latitude. This procedure did not affect the results qualitatively, showing they are robust. Our temporal sampling is centered mainly on two major time points (1997 and 2009/2010), which warrants caution when interpreting fine-scale temporal dynamics. Still, these two points span more than 100 fly generations, equivalent to millennia of human evolution, providing substantial power to detect long-term genomic change. Although additional time points would help refine rate estimates, our data set already captures a robust and biologically meaningful signature of long-term evolutionary change.

If most of the genomic clines were initially the product of secondary contact as previously proposed (Bergland et al. 2016), we would expect samples to become more similar in more recent times and to see fewer clinal variants, which aligns perfectly with our observations. Altogether, this evidence suggests that, as time has passed, gene flow has gradually homogenized the populations along the East Coast, leading to an overall reduction in clinality. Our study highlights the outcome, and fate, of adaptive genetic diversity along a latitudinal gradient, which was dictated by the interplay between natural selection, gene flow, and genomic architecture (i.e., recombination rate and chromosomal inversions). More broadly, our results show that adaptive variation can persist despite substantial gene flow, emphasizing that selection is sufficiently strong to maintain locally favored alleles over decades.

## Methods

### Data Collection

Collection sites and collection information are available in Figure 1 and supplementary Table S1. Most of our historical samples are from previous studies that only focused on a small group of SNPs or allozymes (Sezgin et al. 2004; Lavington et al. 2014). Isofemale lines were founded based on wild-caught flies in the collection sites. The direct offspring of those flies were used to identify *D. melanogaster*. We performed pool-sequencing on our samples using two females from each collected lineage. Direct descendants (F1) of the wild-caught females were kept frozen in alcohol until library preparation. The number of flies per pool varied from 60 to 198 (Table S1). We complemented our sampling points with three pool-seq populations available in NCBI (Project PRJNA256231, see S1) originally from (Bergland et al. 2014). We downloaded the samples using the fastq-dump command from the SRA-toolkit (Sherry et al. 2012). In addition to those samples, one genome sample was donated by Walter Eanes (Veeramah et al. 2020). Because this last sample had an extremely high coverage depth (mean 425X), we performed a pre-downsample step with seqtk (Li 2013), initially reducing the mean depth to 50X.

### Data Cleaning, Mapping, and SNP Calling and Annotation

We used fastp (Chen 2023) to perform the first quality control check and trim adapters when needed. We mapped the sequences to the reference *D. melanogaster* genome r6.25 (Larkin et al. 2021) with bwa (algorithm MEM) version 0.7.15 (Li and Durbin 2010). PCR duplicates were marked with SAMtools version 1.3 (Danecek et al. 2021) as was the conversion from SAM to BAM. Given that the number of flies per pool and the mean sequence depth varied substantially among samples (Table S2) we downsampled the sequencing data so that all samples had a mean effective number of chromosomes (NE) of 20. To determine the sequencing depth (D) required for each sample, we solved the equation:

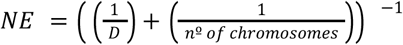

for (D), equating NE to 20, where (n) is the number of chromosomes represented in the pool. Because two female flies were used per isochromosomal lineage and, on average, half of their genomes are identical by descent, we considered each lineage to contribute an average of three chromosomes, corresponding to 1.5 chromosomes per fly. We therefore adjusted the number of chromosomes represented in each pool accordingly, with the exception of pools obtained from (Bergland et al. 2014), for which a single fly per lineage was used. This correction follows the same rationale and effective chromosome estimation approach used in our previous studies based on older methodologies for estimating frequencies of specific SNPs from pooled samples and isofemale line collections (Lavington et al. 2014; Cogni et al. 2015). Because the downloaded sequences were composed of male flies we performed the downsamples separately on the X chromosome of them. Using SAMtools we generated a mpileup with base alignment quality computation (BAQ) enabled, filtering for a base and mapping quality of 20. SNPs were called with PoolSNP (Kapun et al. 2020), with a minimum coverage of 15, minimum alternative allele count across all samples of 5, minimum frequency of 0.001, and missing fraction of 0.1. We used Martin Kapun’s custom Python scripts to filter 5bp up-downstream indels (Kapun et al. 2020). We filtered repetitive regions based on the RepeatMasker library for *D. melanogaster* (obtained from http://www.repeatmasker.org). SNPs were annotated with SnpEff (Cingolani et al. 2012) and we reduced the updownstream definition to 1Kb. For all analyses, we only focused on chromosomes 2, 3, and X. Because of the pooled nature of the data, we did not perform singleton-based quality checks (as heterozygous to non-ref homozygous ratio, or over-representation of A/T nucleotides), instead we computed the transition to transversion ratio based on presence or absence of SNPs per population.

### Nucleotide Diversity, Genetic Differentiation (F_ST_) and Ancestry

We computed Tajima’s pi using PoolGen_var (Kofler et al. 2011; Kapun et al. 2020) on 200kb non-overlapping windows not accounting for missing data. We average window diversity values weighted by the true number of sites in each window to compute the genomic diversity values. To understand if diversity changed across the years we ran a mixed linear model of the mean diversity of each population against its collection year and latitude (with no interaction), considering the random effect of the chromosome on the intercept. We then computed global F_ST_ between all pairs of populations using the autosomes or the X chromosome with grenedalf (Czech et al. 2024), using the Hudson estimator (Hudson et al. 1992) and filtering for missing data. To look at patterns of isolation by distance we regressed the pairwise F_ST_ to the geographic distance between populations. This was made separately to samples of 1997 and 2009/2010. Because CMD97B had a smaller coverage depth and was more differentiated from other samples, we also conducted these analyses without it.

Principal component analysis (PCA) was made only with variants with depth ≥ 20X after filtering for missing data using the prcomp function with custom R (R Core Team 2022) scripts, as was the PC’s regression with environmental variables. Environmental variables of the prior three months before collection dates were obtained with the nasapower R package (Sparks 2018).

To compute F_ST_ with ancestral populations and ancestry proportions as described by Bergland et al (2016), we assembled ancestral reference panels (Table S8) composed of 23 European and 23 Western-Central African haploid genomes. European genomes were from France and Sweden (project PRJNA30085 and PRJNA294138) (Pool et al. 2012; Kapopoulou et al. 2020). African genomes were from Western-Central Africa (Cameroon, Gabon, Nigeria, and Guinea) (project PRJNA30085). Given the European introgression on African populations, we only used genomes with less than 11% introgression, to minimize this confounding factor. We used the same pipeline we used with pool-sequencing to map those genomes but used the GATK v4.3.0.0 (McKenna et al. 2010) variant calling pipeline to call SNPs. Briefly, we constructed gvcf files with GATK haplotypecaller setting the ploidy to 1, then assembled a genomic database with GenomicsDBImport and called variants with GenotypeGVCFs. We then removed indels and repetitive regions. We constructed a sync file combining the ancestral panel with our samples, treating the European and African sets as pools of 23 individuals and 23 read depth, in which each individual’s allele would count as one read. To compute F_ST_ between our samples and this panel we also used grenedalf (Czech et al. 2024). We ensured that all positions that were polymorphic in at least one dataset (the ancestral panel or our samples) and had enough quality in both datasets were included. In other words, we ensured that positions that were polymorphic in one dataset but fixed for the reference in the other were comprehended. However, we applied a 0.1 MAF filter as it is known that rare variants can influence F_ST_ results (Bhatia et al. 2013). To compute ancestry proportions we sampled 5000 SNPs and performed an intercept-free regression model of those SNPs against the frequency of the SNPs in the African and European populations. We repeated this regression 100 times and used the mean of the coefficients as the proportion of African and European ancestry of each population. We restricted this last analysis to informative polymorphisms, SNPs with at least a 10% frequency difference between African and European populations.

In addition to those methods, we also used f4 ratio to compute ancestry proportions. To do this, we assembled another ancestral panel, composed of the same samples used in the first plus 64 haploid Zambian (project SRP006733) (Lack et al. 2015) samples and 32 inbred Egyptian samples (project PRJNA327349) (Lack et al. 2016). We used the same mapping and calling strategy as the first one and also constructed a sync file combining this ancestral panel with our samples. For that, we considered each ancestral population as a pool with maximum depth of the total number of genomes used. Because the Egyptian lines still had some heterozygous regions, whenever a region was polymorphic, we sampled only one allele to be included. After constructing the sync files, we computed the f4 ratio with poolfstat R package (Gautier et al. 2022), using a MAF filter of 0.05 and a minimum coverage per pool of 10X. For the numerator of the f4 ratio, we used F4(EG,ZI;westAFR,x) and for the denumerator we used F4(EG,ZI;westAFR,EU), where EG, ZI, EU and westAFR are the Egyptian, Zimbabwean, European and west African populations respectively and x is the focal North American population.

### Identification and Enrichment of Clinal SNPs

To identify clinal SNPs we used a binomial linear model (glm) of allele frequency against latitude for each SNP. This was done separately for 1997 and 2009/2010, we only used those periods in this analysis because we did not have enough populations in 2017 or 2022/2023. Moreover, we only considered biallelic SNPs and SNPs that were present (polimorphic) in at least three populations in the analyzed time points, and which site passed the quality filters for sequencing in at least 5 populations. The models were weighted for the effective number of chromosomes, accounting for sequence depth and the number of flies used in each pool (Kolaczkowski et al. 2011; Bergland et al. 2014; Rodrigues et al. 2021). To define clinal SNPs we computed False Discovery Rates (FDR) (Storey and Tibshirani 2003) with R package qvalue (Storey at al. 2022). To run this analysis, we used two different sets of populations: using all populations available in both periods; and, using only populations that matched both periods (MFL97 and RVA97 removed from 1997 and JFL10, dlGA10, and dlSN10 removed from 2009/2010). We used R to perform a Kolmogorov-Smirnov and a McNemar’s Chi-squared test to compare the p-values distribution of both periods and the stability of clinal SNPs across time.

To assess the power of our clinal analysis under the observed sampling and sequencing conditions, we performed simulations using the empirical metadata for each sampling period (1997 and 2009/2010), including latitude, NE, and sequencing depth for each population and SNP. We simulated a mixture of clinal and non-clinal SNPs by assigning 75% of SNPs a slope of zero and the remaining 25% slopes evenly distributed between 0.05 and 1. For each SNP, expected allele frequencies were generated from a logistic relationship with standardized latitude, with the intercept fixed at zero. We then simulated allele counts using the observed NE and, subsequently, pooled sequencing read counts using the observed sequencing depth. The resulting pooled allele frequencies were analyzed using the same binomial generalized linear model as the empirical data, with latitude as the predictor and NE as the model weight. For each simulated SNP, we recorded the estimated slope and associated P-value, and repeated the procedure for 100 simulation replicates, randomly reassigning slopes to SNPs in each replicate.

To check the agreement between clinal variants, we first measured the proportion of clinal SNPs that were clinal in both periods relative to 1997 or 2009/2010. We then measured the proportion of clinal SNPs that had the same slope sign in both periods. Finally, focusing on concordant clinal SNPs, we assessed whether the strength of latitudinal clines changed between sampling periods by performing a deming regression analysis of slope coefficients in 1997 and 2009/2010. We estimated the regression intercept and slope and their 95% confidence intervals. We additionally tested whether the regression slope differed from 1, which would indicate a systematic change in cline strength between periods, using a two-sided normal test based on the estimated slope and its standard error.

To verify if recombination rate had influence over presence of clinal SNPs we downloaded the recombination map from (Comeron et al. 2012). We then build general mixed binomial models with the R package lme4 (Bates et al. 2015) using the clinality of a SNP (1 with if it is clinal under a 10% FDR and 0 if not) as response variable and the recombination rate as predictor with the random effect of the genomic window in which recombination was calculated. To verify if recombination rate had an effect on the lost of clinal SNPs, we only used SNPs that were clinal under 10% FDR in 1997, we then used the same general model, with the persistence of a SNP as clinal (1 if it is no longer clinal in 2009/2010 and 0 if it remains clinal in 2009/2010) as a response variable.

To look for enriched genomic regions among clinal SNPs of a specific period, we used the annotations done by SnpEff (Cingolani et al. 2012). Since each SNP can have more than one functional annotation and because SnpEff sorts higher quantitative effects first, we used only the first annotation. We simplified annotations in one of the following categories: intergenic, intronic, 1Kb upstream, 1Kb downstream, 5’UTR, 3’UTR, splice, synonymous coding, and nonsynonymous coding (Table S21). We then computed the odds ratio between the change of a clinal SNP being in one of those regions to that of 100,000 randomly sampled non-clinal SNPs. We repeated the sampling of the random SNPs 10,000 times and recomputed the odds ratio each time.

### Signature of selection and GO enrichment terms

To look for selection signatures we computed window F_ST_ between pairs of samples from the same location, but different time points. To do that we also used grenedalf (Czech et al. 2024) and the Hudson estimator (Hudson et al. 1992). Windows were 100 SNPs in length striding 10 SNPs in each new window. We then used the top 0.1% outlier windows to search for possible selected regions. Adjacent top windows within 4 kb of each other were merged using BEDTools v2.30.0 (Quinlan and Hall 2010), and candidate overlapping genes within these regions were obtained using the *D. melanogaster* GTF 6.55 file (Öztürk-Çolak et al. 2024) and a custom python script.

To find enriched GO terms we associated each unique gene with its corresponding GO categories. Then we counted how often each GO term occurred among outlier genes versus the full gene set. We repeated this count permuting the location of the outlier regions along the genome, preserving the observed distribution of outlier window sizes and chromosomes. We repeated the permutation step 500 times and used it to compute the p-values based on the empirical probability that a GO term would appear as often (or more often) in the permuted regions than in the real data. We repeated this process 20 times and averaged the p-values of each GO term.

### Data availability

Raw short-read sequence data generated in this study are available in the NCBI Sequence Read Archive under the project number PRJNA1415801.

### Code availability

All code used in the analysis of this study is available on GitHub (https://github.com/VitoriaHorvathMiranda/time_clines_workflow).

## Acknowledgements

We thank Cristina Yumi Miyaki, Maria Dulcetti Vibranovski, Débora Yoshihara Caldeira Brandt, Julián Mensch and members of the Ecologia Evolutiva lab for reading and providing comments on this manuscript, and Brian C. Verrelli for collecting the 1997 samples. This study was funded by Coordenação de Aperfeiçoamento de Pessoal de Nível Superior (CAPES; n° 88887.683802/2022-00) and the São Paulo Research Foundation (FAPESP; processes 2013/25991-0, 2022/12549-6, 2023/09079-0, 2021/06874-9), a Harvard University Lemann grant, and a Newton Advanced Fellowship from the Royal Society.

## Notes

### Competing Interest Statement

The authors have declared no competing interest.

### Summary of Updates

In brief, we improved our statistical analyses (tackling the differences in power across years) and expanded our sensitivity analysis. We better contextualized our work with earlier cytogenetic clinal studies and more recent genomic resources. We have also clarified our description of sample collection and storage. Further, we tightened the narrative and streamlined the supplemental material.

